# Closure motif-mediated state changes of the HORMA protein ASY1

**DOI:** 10.1101/2020.05.04.064501

**Authors:** Chao Yang, Bingyan Hu, Stephan Michael Portheine, Pichaporn Chuenban, Arp Schnittger

## Abstract

Meiotic HORMA domain-containing proteins (HORMADs) are key components of the chromosome axis and play an essential role in meiosis in many organisms. The meiotic HORMADs, including yeast Hop1, mouse HORMAD1 and HORMAD2, and Arabidopsis ASY1, assemble along chromosomes at early prophase and the closure motif at their C-termini has been hypothesized to be instrumental for this step by promoting HORMAD oligomerization. In late prophase, ASY1 and its homologs are progressively removed from synapsed chromosomes promoting chromosome synapsis and recombination. The conserved AAA+ ATPase PCH2/TRIP13 has been intensively studied for its role in removing HORMADs from synapsed chromosomes. In contrast, not much is known about how HORMADs are loaded onto chromosomes. Here, we reveal that the PCH2-mediated dissociation of the HORMA domain of ASY1 from its closure motif is important for the nuclear targeting and subsequent chromosomal loading of ASY1. This indicates that the promotion of ASY1 to an ‘unlocked’ state is a prerequisite for its nuclear localization and chromosomal assembly. Likewise, we find that the closure motif is also necessary for the removal of ASY1 by PCH2 later in prophase. Our work results in a grand unified model for PCH2 and HORMADs function in meiosis.

## Introduction

A fundamental process for sexual reproduction is meiosis during which one round of DNA replication is followed by two consecutive nuclear divisions resulting in the reduction of the chromosome set of a cell by half. Importantly, genetic information from maternal and paternal progenitors is re-shuffled during meiosis through homologous recombination, thereby generating genetic variation and thus contributing to the diversity of life.

Recombination requires the formation of a meiosis-specific proteinaceous structure called chromosome axis that spans along the entire length of the chromosomes. The chromosome axis is believed to organize sister chromatids into a loop-array configuration and thus facilitates interactions of homologous chromosomes (1–3). The chromosome axis consists of cohesin complexes encompassing sister chromatids, and other meiosis-specific proteins including the meiotic HORMA domain-containing protein (HORMADs) family (ASY1 in Arabidopsis, Hop1 in yeast; HORMAD1/2 in mouse); the coiled-coil domain-containing ‘linker’ proteins (ASY3 in Arabidopsis, Red1 in yeast; SYCP2 in mouse); and the small coiled-coil protein family proteins (ASY4 in Arabidopsis, the mammalian SYCP3/SCP3 homolog) (4–11). Current models suggest that the coiled-coil proteins form filamentous complexes that form the core of the chromosome axis. These filaments are thought to localize on chromosomes through binding to cohesin complexes and thereby organize the loop-array structure of meiotic chromosomes (12).

The meiotic HORMADs are involved in many key meiotic events, i.e. double-strand break (DSB) formation, synaptonemal complex assembly and crossover formation (4, 5, 13, 14). These proteins are characterized by an N-terminal HORMA domain (for <u>Hop</u>1, <u>R</u>ev7 and <u>MA</u>D2), a conserved domain which interacts with a short sequence motif termed ‘closure motif’ (15, 16). The HORMA domain-to-closure motif interaction is thought to anchor ASY1 and other HORMADs to the axis via interaction with the closure motif of ASY3 and its orthologs (9, 12, 17).

However, all meiotic HORMADs studied, including Hop1, HORMAD1/2 and ASY1 also contain a closure motif at their own C-terminus, and because of this, these HORMADs tend to fold into a HORMA domain-closure motif bound state called self-’closed’ state (12, 15, 17, 18). This self-closed state of HORMADs is expected to block the binding of their HORMA domain with the closure motifs of the linker proteins such as ASY3, raising the question of how this interaction is formed *in vivo* (15, 17, 18). It has been proposed that the conformational conversion from a self-closed to the ‘unlocked’ state might be mediated by the conserved AAA+ ATPase PCH2/TRIP13 protein (15, 17, 19). However, there is no experimental evidence to support this up to now.

The meiotic HORMADs undergo a highly dynamic assembly and disassembly process, which is essential for the homologous chromosome synapsis and recombination. The removal of meiotic HORMADs from chromosome axes at late prophase is catalyzed by PCH2/TRIP13 and has been extensively studied in different organisms including budding yeast, mouse and Arabidopsis (Lambing et al, 2015; Joshi et al, 2009; Wojtasz et al, 2009). However, how the chromosomal assembly of meiotic HORMADs is controlled is much less clear. Recently, we have shown that PCH2 is also implicated in nuclear targeting of ASY1 during early prophase (18). However, nothing is currently known about how PCH2 controls this process.

Here, we present the mechanism of the PCH2-regulated nuclear targeting and chromosome localization dynamics of ASY1. We discovered that PCH2 regulates the nuclear targeting of ASY1 by preventing the binding of the HORMA domain with the closure motif. Thus, ASY1 is required to be in an unlocked state for its nuclear import. Moreover, we revealed that the position and origin (the sequence *per se*) of the closure motif are not essential for its functional interplay with PCH2.

## Results

### The closure motif is not required for chromosomal association of ASY1

Previous experiments suggested that the closure motif is involved in the chromosome association of ASY1 (17, 18, 20). To test this, we introduced the previously generated deletion construct of ASY1 (*PRO_ASY1_:ASY1^1-570:GFP^*, designated *ASY1^Δclosure^:GFP*) in which the closure motif comprising the last 25 amino acids (aa) of ASY1 (596 aa in total) was removed from a functional reporter construct (*PRO_ASY1_:ASY1:GFP*, called *ASY1:GFP*) (18), into *asy1* mutants (Fig. 1A).

**Figure 1.**
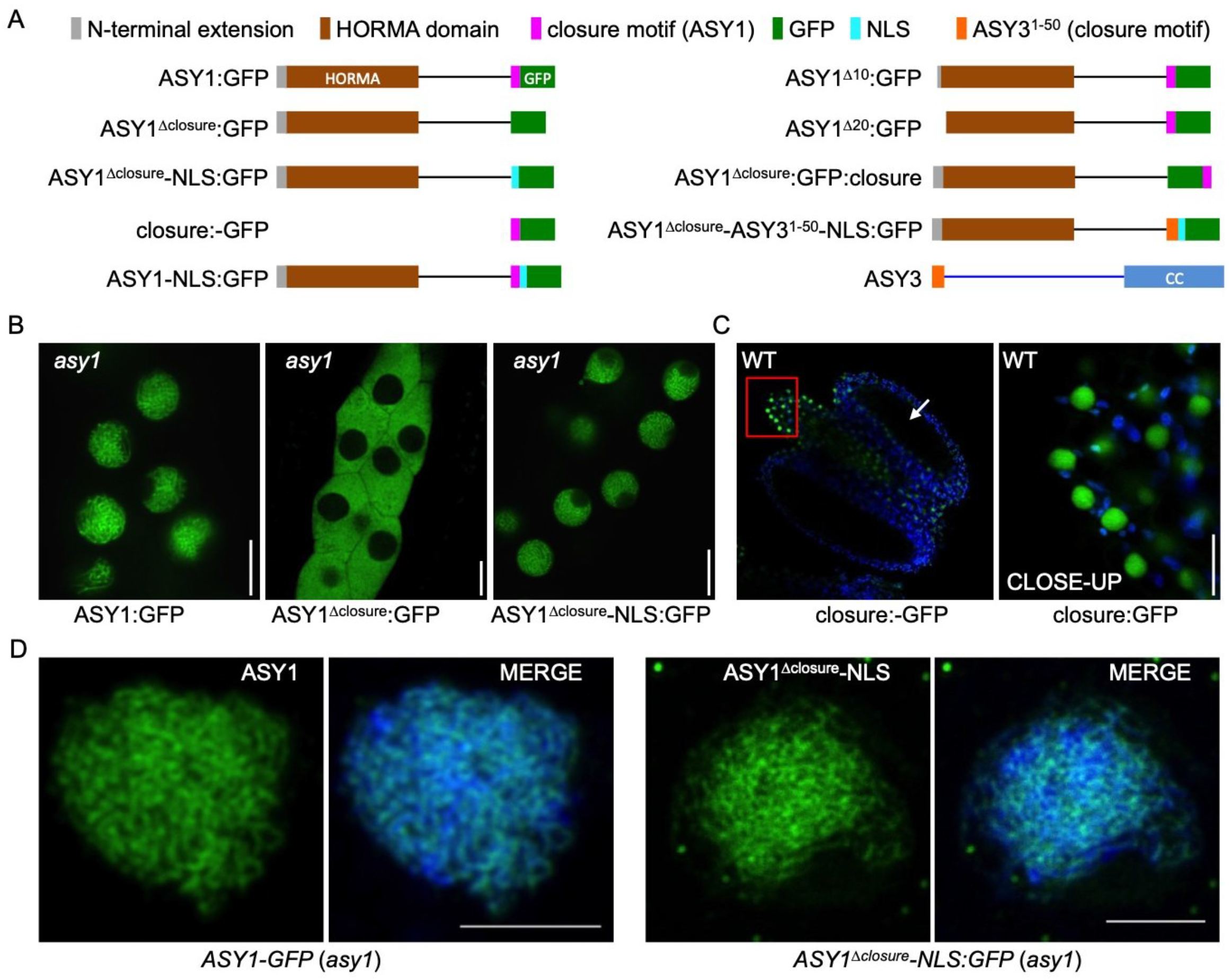
Closure motif of ASY1 functions as a nuclear localization signal (NLS) and is not required for its chromosomal localization. (A) Schematic of different ASY1 versions and ASY3 used in this study. (B) Localization of ASY1:GFP, ASY1^Δclosure^:GFP, and ASY1^Δclosure^-NLS:GFP in the male meiocytes of *asy1* mutants using confocal microscopy. Bar: 10 μm. (C) Localization of closure:GFP in the anther of wildtype using confocal microscopy. The CLOSE-UP shows the magnification of the connective tissue highlighted by the red rectangle. Bar: 10 μm. (D) Immunolocalization of ASY1:GFP and ASY1^Δclosure^-NLS:GFP at early prophase in male meiocytes of *asy1* mutants using antibody against GFP. MERGE shows the overlay of ASY1 signal and DAPI-stained DNA. Bar: 5 μm.

In *asy1* mutants harboring the full length reporter, ASY1:GFP accumulated in the nuclei of male meiocytes during meiotic prophase I, as reported previously (18) (Fig. 1B, Table 1). In contrast, ASY1^Δclosure^:GFP was found to be present only in the cytoplasm and no signal was detected in the nucleus, suggesting that the closure motif plays a role in nuclear targeting of ASY1 (Fig. 1B). To test this possibility, we generated plants containing a construct, in which only the closure motif sequence fused with GFP driven by the *ASY1* promoter (*PRO_ASY1_:ASY1^571-596^:GFP*, called *closure:GFP*) was present (Fig. 1A). Unexpectedly, we did not detect any signal of the closure:GFP in male meiocytes indicating that for the proper expression in meiocytes sequence information from the introns of *ASY1* are necessary (Fig. 1C, arrow). Nonetheless, closure:GFP was found to be expressed in the epidermal cells of the connective tissue of anthers, where it specially localized in nuclei, corroborating that the closure motif of ASY1 functions as a nuclear localization signal (NLS) (Fig. 1C, CLOSE-UP).

**Table 1.**
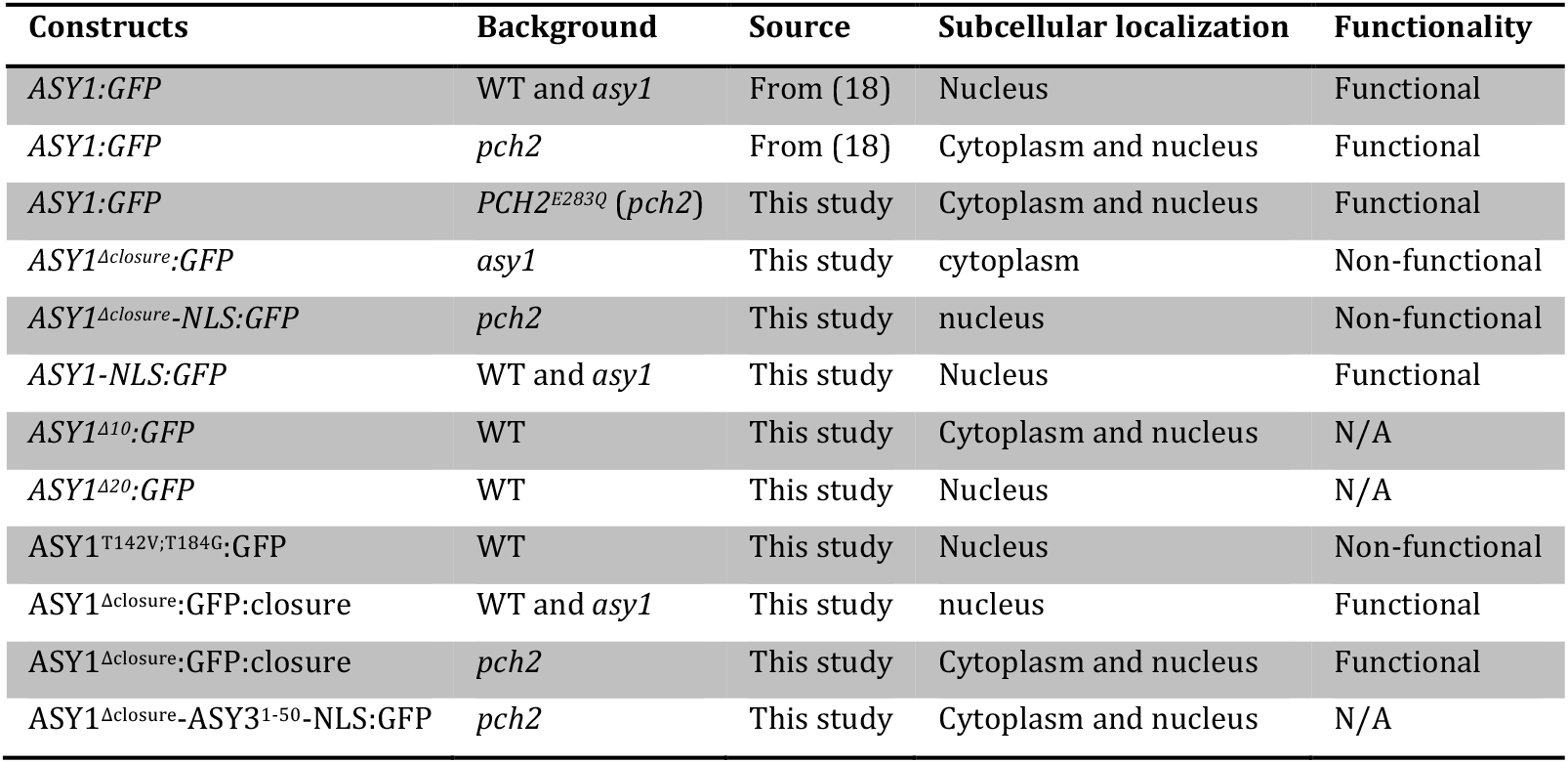
Summary of the subcellular localization of different versions of ASY1 used in this study in different plant backgrounds. N/A denotes Not Analyzed.

To check whether the closure motif is required for chromosomal association of ASY1 after nuclear import, we next generated a separation-of-function allele, in which the closure motif was substituted with the NLS sequence of the SV40 Large T-Antigen (PKKKRKV) (*PRO_ASY1_:ASY1^1-570^-NLS:GFP*, called *ASY1^Δclosure^-NLS:GFP*) (Fig. 1A). Notably, ASY1^Δclosure^-NLS:GFP was exclusively present in the nuclei of male meiocytes in *asy1* mutants suggesting that the closure motif indeed functions in nuclear targeting of ASY1 (Fig. 1B, Table 1).

To further dissect the importance of the closure motif for the function of ASY1, we first performed immunofluorescence experiments using an antibody against GFP and compared in detail the localization of ASY1:GFP and ASY1^Δclosure^-NLS:GFP in male meiocytes of *asy1* mutants. To this end, we found that ASY1^Δclosure^-NLS:GFP showed an indistinguishable chromosomal association from ASY1:GFP forming a thread-like signal along all chromosomes at early prophase (Fig. 1D). This was also confirmed by the localization patterns of ASY1:GFP and ASY1^Δclosure^-NLS:GFP in live cells of *asy1* mutants by confocal microscopy (Fig. S1A). Taken together, these results suggest that the closure motif of ASY1 is not essential for its chromosomal association.

### Deletion of the closure motif dominantly interferes with synapsis

Given the apparently correct localization on chromosomes, we asked whether ASY1^Δclosure^-NLS:GFP was also functional. However, while ASY1:GFP fully rescued the fertility defects of *asy1* mutants, ASY1^Δclosure^-NLS:GFP did not rescue the *asy1* mutants, as revealed by short siliques, high pollen abortion and defective chromosome segregation (Fig. S2). This suggests that ASY1 without the closure motif, despite its proper localization, is not functional. Notably, ASY1^Δclosure^-NLS:GFP functioned in a dominant negative manner since we also observed apparent fertility defects in 27 out of 35 T1 plants when this construct was introduced into a wild-type background (called *ASY1^Δclosure^-NLS:GFP*/WT) (Fig. S2A and B).

To exclude that this loss of ASY1 functionality was due to the added NLS signal, we created a full-length version of ASY1 with this NLS sequence (*PRO_ASY1_:ASY1-NLS:GFP*, called *ASY1-NLS:GFP*) (Fig. 1A) and introduced this construct into wild-type and *asy1* mutant plants. Most of the wild-type T1 transformants (35 out of 40) expressing the ASY1-NLS:GFP (called *ASY1-NLS:GFP*/WT) were fully fertile, and the fertility of *asy1* mutants harboring this construct was fully restored (Fig. S2A and B), suggesting that ASY1-NLS:GFP was a functional protein.

Next, we compared the *ASY1-NLS:GFP*/WT and the *ASY1^Δclosure^-NLS:GFP*/WT plants in detail. Both ASY1 variants showed a tight chromosomal association at early stage of meiotic prophase I (Fig. S1B). Subsequent chromosome spread analyses revealed that, consistent with the high level of fertility, *ASY1-NLS:GFP*/WT plants followed a regular meiotic path in which five paired bivalents were aligned on the metaphase I plate and then segregated equally at the first and second meiotic divisions, resulting in the formation of daughter cells with an equal number of chromosomes in the tetrad stage (Fig. 2A). While no obvious differences were observed until zygotene compared to the *ASY1-NLS:GFP*/WT plants, no meiocytes at pachytene stage were found in *ASY1^Δclosure^-NLS:GFP*/WT plants (n=98) (Fig. 2A). Instead, only a partial synapsis was observed in *ASY1^Δclosure^-NLS:GFP*/WT plants. A defective synapsis in *ASY1^Δclosure^-NLS:GFP*/WT plants was further confirmed by the detection of ZYP1 in immunofluorescence experiments (Fig. 2B). Defects in synapsis are likely the reason for the frequent observation of univalents at metaphase I (76%, n=17) followed by unbalanced chromosome segregation at the first and second meiotic division (Fig. 2A). This is likely the reason for the high level of pollen abortion and reduced fertility (Fig. S2A and B). These results suggest that ASY1 without the closure motif dominantly interferes with the progression of synapsis.

**Figure 2.**
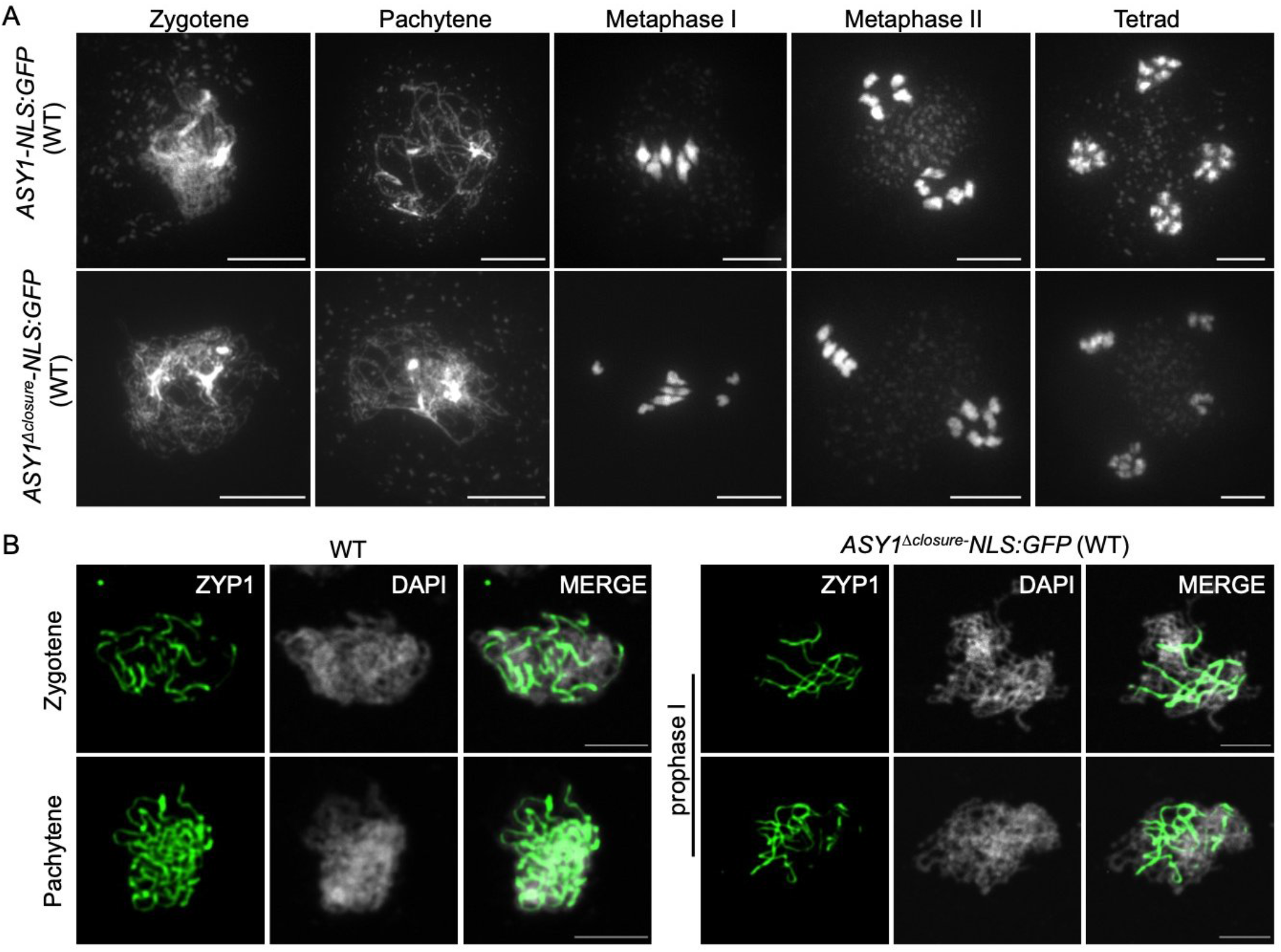
Deletion of the closure motif dominantly interferes with synapsis. (A) Chromosome spread analysis of male meiocytes at different meiotic stages in wild-type (WT) plants harboring *ASY1-NLS:GFP* or *ASY1^Δclosure^:NLS-GFP* construct. Bar: 10 μm. (B) Immunolocalization of ZYP1 at different meiotic prophase I stages in wild-type plants harboring *ASY1-NLS:GFP* or *ASY1^Δclosure^:NLS-GFP* construct. Bar: 5 μm.

### The closure motif is required for the removal of ASY1 from chromosomes

The defective synapsis in *ASY1^Δclosure^-NLS:GFP*/WT plants together with the tight association of ASY1^Δclosure^-NLS:GFP with the chromosomes at late prophase I (Fig. S1B), is reminiscent of the phenotype of *pch2* mutants in which the removal of ASY1 from chromosomes is compromised resulting in synapsis defects (21). Thus, we wondered whether the deletion of the closure motif might lead to a defective removal of ASY1^Δclosure^-NLS:GFP from synapsed chromosomes and hinder the progression of synapsis.

To answer this, we checked the presence of ASY1^Δclosure^-NLS:GFP at synapsed chromosomes in male meiocytes of wildtype by the co-immunolocalization analysis of ASY1 and ZYP1 using antibodies against GFP and ZYP1. The *ASY1-NLS:GFP*/WT plants were used as a control. Indeed, while ASY1-NLS:GFP in the wild-type background was removed from the synapsed regions like normal ASY1:GFP (Fig. 3A), the ASY1^Δclosure^-NLS:GFP signal was still not reduced at late prophase I, and its signal intensity at synapsed chromosomes appeared even stronger than that in non-synapsed regions (Fig. 3B). This suggests that the presence of the closure motif is required for the efficient depletion of ASY1 when chromosomes become synapsed.

**Figure 3.**
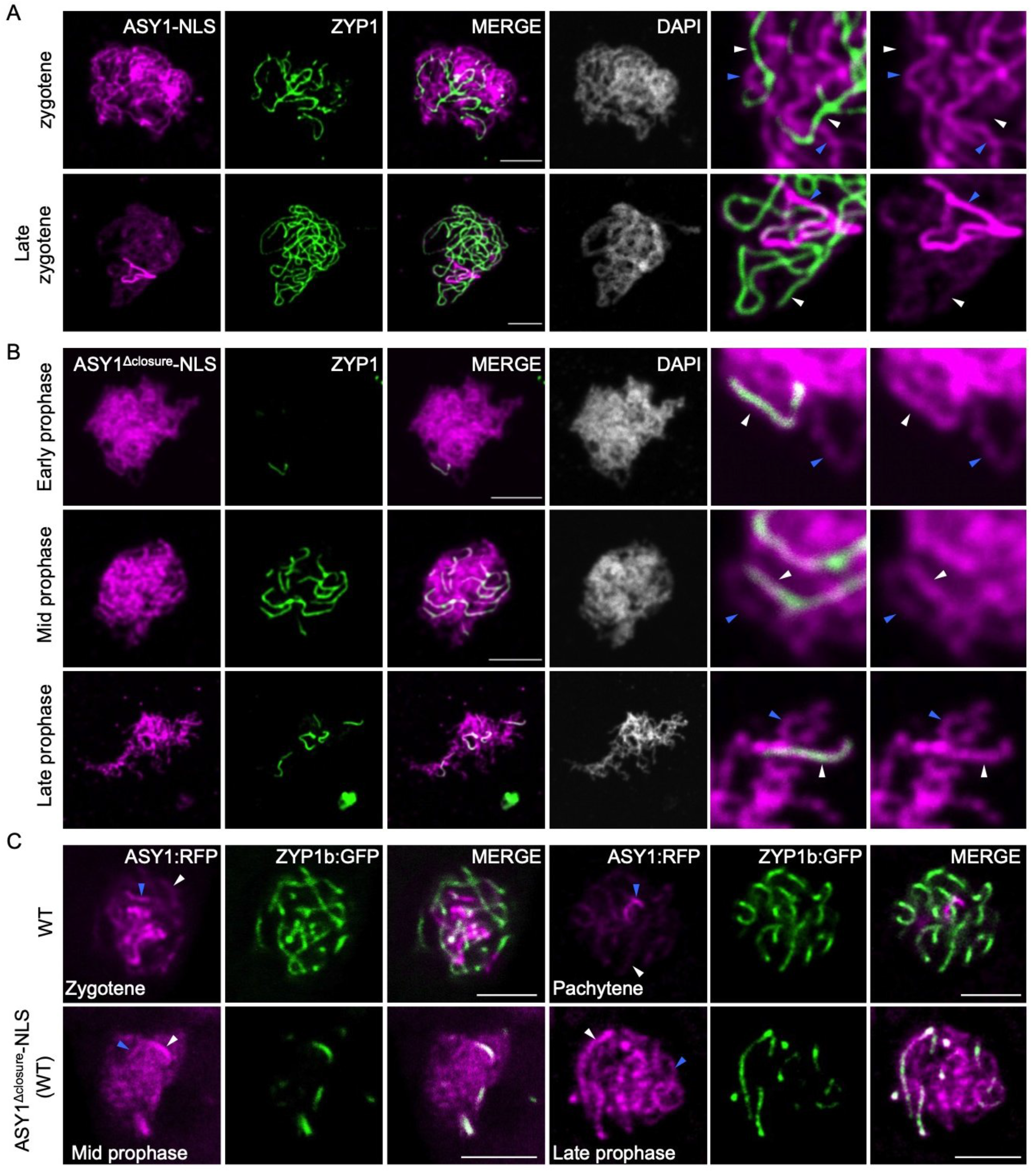
The closure motif is required for the PCH2-mediated removal of ASY1 from synapsed chromosomes. (A and B) Co-immunolocalization of ASY1-NLS:GFP (A) or ASY1^Δclosure^-NLS:GFP (B) with ZYP1 in male meiocytes of wildtype plants using antibody against GFP and ZYP1. (C) Co-localization of ASY1:RFP with ZYP1b:GFP in male meiocytes of wildtype (WT) and *ASY1^Δclosure^-NLS*/WT plants using confocal microscopy. All blue and white arrows indicate the non-synapsed and synapsed regions, respectively. Bar: 5 μm.

The defective synapsis in *ASY1-NLS:GFP*/WT plants might also be due to a competition between a non-functional ASY1^Δclosure^-NLS:GFP and the wild-type version of ASY1 for chromosomal binding in a non-preferential manner that would reduce the abundance of the functional ASY1 on chromosomes. If this were the case, the chromosomal removal of the wild-type version of ASY1 would also be likely affected by the presence of ASY1^Δclosure^-NLS:GFP. To test this hypothesis, we introduced *ASY1^Δclosure^-NLS* without the GFP tag into wild-type plants expressing a full-length version of the ASY1 functional reporter (*PRO_ASY1_:ASY1:RFP*, called *ASY1:RFP*) together with a ZYP1b reporter (*PRO_ZYP1b_:ZYP1b:GFP*, called *ZYP1b:GFP*), generated previously (18, 22). Compared to the control plants without *ASY1^Δclosure^-NLS*, ASY1:RFP appeared to persist at synapsed regions in the plants expressing *ASY1^Δclosure^-NLS*, and the signal intensity of ASY1:RFP at synapsed chromosomes appeared even stronger than that in the non-synapsed parts reflecting the presence of ASY1^Δclosure^-NLS:GFP in the wildtype (Fig. 3B and C). Thus, the non-functional ASY1^Δclosure^-NLS:GFP and the wild-type ASY1 version form very likely a mixed layer along the chromosome axis, interfering with chromosome synapsis.

Taken together, we conclude that the closure motif of ASY1 is required for its efficient removal from the synapsed chromosomes.

### An interplay between the closure motif and PCH2 controls the nuclear targeting and chromosomal association of ASY1

Previously, we have reported that PCH2 is essential for the efficient nuclear targeting of ASY1, as seen by the accumulation of ASY1:GFP in the cytoplasm of male meiocytes of *pch2* mutants (Fig. 4A) (18). In contrast, we found that ASY1^Δclosure^-NLS:GFP in *pch2* mutants localized exclusively to the nuclei of male meiocytes (Fig. 4B, Table 1). To explore the epistatic relationship between the closure motif and PCH2 for the nuclear targeting of ASY1, we introduced the ASY1-NLS:GFP into *pch2* mutants creating a situation in which we had two opposing forces: the promotion of nuclear entry by the NLS versus the cytoplasmic retention of the full-length ASY1 protein caused by the absence of *PCH2*.

**Figure 4.**
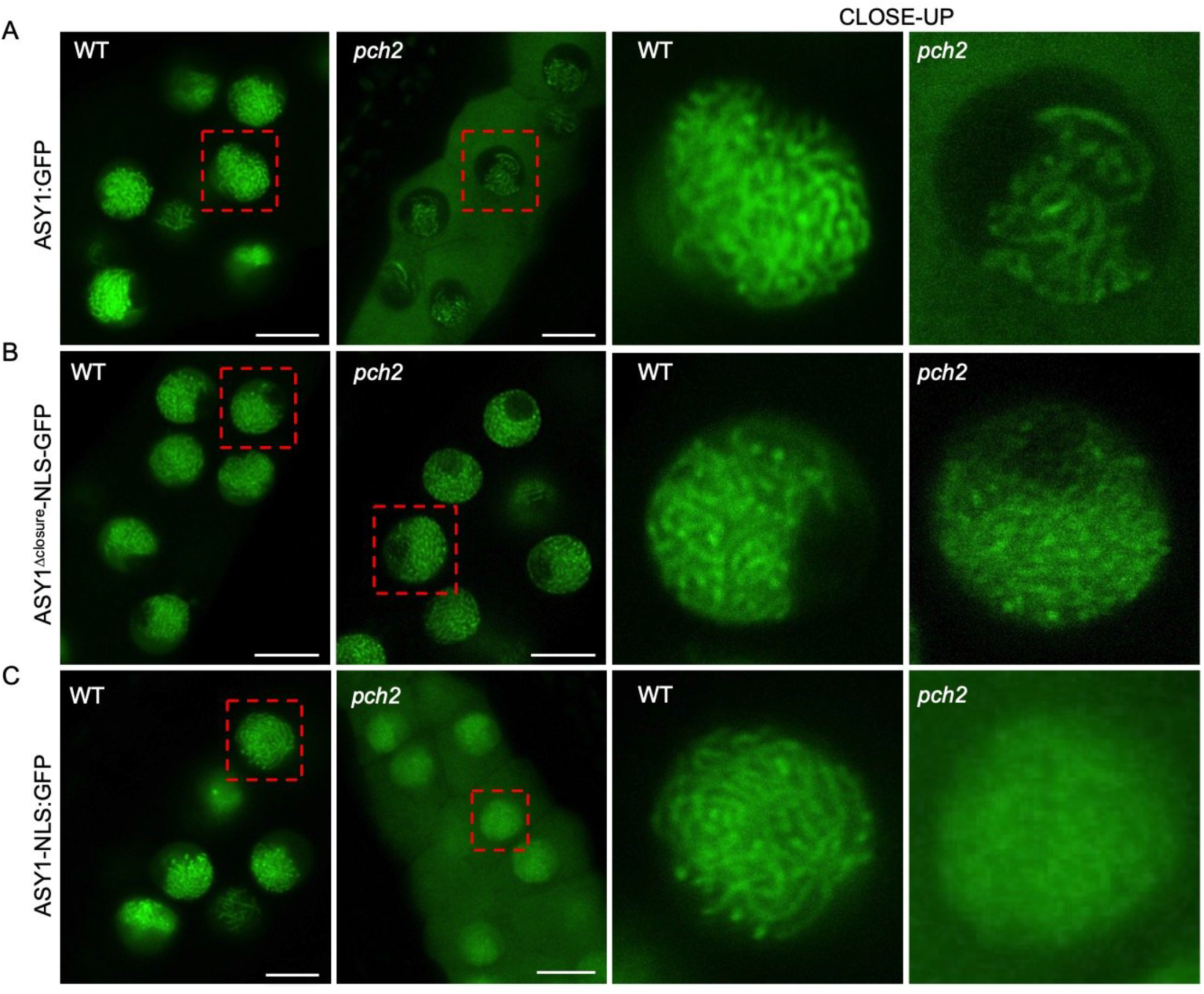
An interplay between the closure motif and PCH2 controls the nuclear targeting and chromosomal association of ASY1. Localization of ASY1-GFP (A), ASY1^Δclosure^-NLS:GFP (B), and ASY1-NLS:GFP (C) in male meiocytes of wildtype and *pch2* mutants. Bar: 10 μm.

We found a higher level of ASY1-NLS:GFP present in the nucleus compared to that of ASY1:GFP in *pch2* mutants (Fig. 4A and C), indicating that the NLS sequence used could facilitate the import of a small part of ASY1-NLS:GFP into the nucleus in the *pch2* mutant background. However, we realized that the addition of the NLS cannot fully rescue the nuclear targeting of the full-length ASY1 version in *pch2* mutants and a considerable amount of ASY1-NLS:GFP was still detected in the cytoplasm of male meiocytes (Fig. 4C). Thus, the presence of the NLS sequence is not sufficient to restore full nuclear localization of ASY1 in *pch2* mutants.

At the same time, we noticed that ASY1-NLS:GFP showed a diffuse presence in the whole nucleus in *pch2* mutants at early prophase while it was normally assembled on chromosomes in the wildtype (Fig. 4C). Therefore, we reasoned that the diffuse appearance of ASY1-NLS:GFP in the nucleus of *pch2* mutants is due to the small portion of ASY1-NLS:GFP proteins that are imported into the nucleus aided by the NLS, but cannot properly localize onto the chromosomes as PCH2 is absent. Thus, we conclude that in addition to the well-known role of PCH2 for ASY1 removal at late prophase, it is also indispensable for the normal chromosomal assembly of ASY1 at early meiosis. Notably, the ASY1-NLS:GFP proteins without the closure motif, i.e. ASY1^Δclosure^-NLS:GFP, displayed a clear chromosomal association in *pch2* mutants (Fig. 4B).

Taken together, these results suggest that the nuclear targeting defects of ASY1 in *pch2* mutants result from the presence of the closure motif, i.e. that there is a functional interplay between the closure motif and PCH2 for the efficient nuclear localization of ASY1. In addition, a PCH2-closure motif interaction seems relevant for the chromosomal association of ASY1 at early prophase.

### PCH2 controls the nuclear targeting of ASY1 through regulating the HORMA domain-to-closure motif interaction

PCH2 is thought to antagonize the chromosomal association of the meiotic HORMADs, including ASY1, by driving a conformational conversion of the HORMAD proteins and regulating their interaction with the binding partners, i.e. ASY3 in Arabidopsis (15, 17, 19). In addition, we previously found that ASY1, at least when being expressed in yeast cells, tends to fold into a self-closed state through binding of its closure motif to its HORMA domain (18). These observations, together with the restoration of the nuclear localization of ASY1 in *pch2* mutants by deleting the closure motif (Fig. 4B, Table 1), led us to hypothesize that the interaction of the HORMA domain with the closure motif might account for the cytoplasmic retention of ASY1 when PCH2 is absent. Thus, the efficient nuclear targeting of ASY1 might require the dissociation of the HORMA domain from the closure motif (a conformation called ‘unlocked’ state in this study) mediated by PCH2.

To test this hypothesis, we first analyzed whether the PCH2-mediated conformational change of ASY1 is necessary for its efficient nuclear targeting. The disordered N-termini of meiotic HORMADs have been found to make transient contact with PCH2, an interaction by which PCH2 partially unfolds the protein to allow for the disengagement of the HORMA domain from its interacting sequence, i.e. the closure motif (17, 19, 23). To this end, we generated a reporter version of ASY1 lacking the first 2-10 aa (*PRO_ASY1_:ASY1^Δ2-10:GFP^*, called *ASY1^Δ10^:GFP*) (Fig. 1A), which theoretically should disturb the interaction of PCH2 with ASY1, and then checked the localization of ASY1^Δ10^:GFP in the wild-type background in the presence of PCH2. In support of the idea that the N-terminus of ASY1 is the action site of PCH2, the localization pattern of ASY1^Δ10^:GFP at different prophase I stages in the wildtype phenocopied that of ASY1:GFP in *pch2* mutants (Fig. 4A and 5A, Table 1). This is consistent with the conclusion that the PCH2-mediated conformational change of ASY1 is required for its efficient nuclear targeting. Moreover, we constructed a mutant version of PCH2 in which the glutamic acid (E283) of the conserved Walker B motif involved in ATP hydrolysis (24, 25), was substituted with a glutamine (called *PCH2^E283Q^*) in a functional genomic construct of *PCH2* generated previously (18). Notably, we found that *PCH2^E283Q^* introduced into *pch2* mutants could not rescue the cytoplasmic retention of ASY1:GFP (Fig. S3), indicating that the ATP hydrolyzing activity of PCH2 is required for the conformational conversion of ASY1.

**Figure 5.**
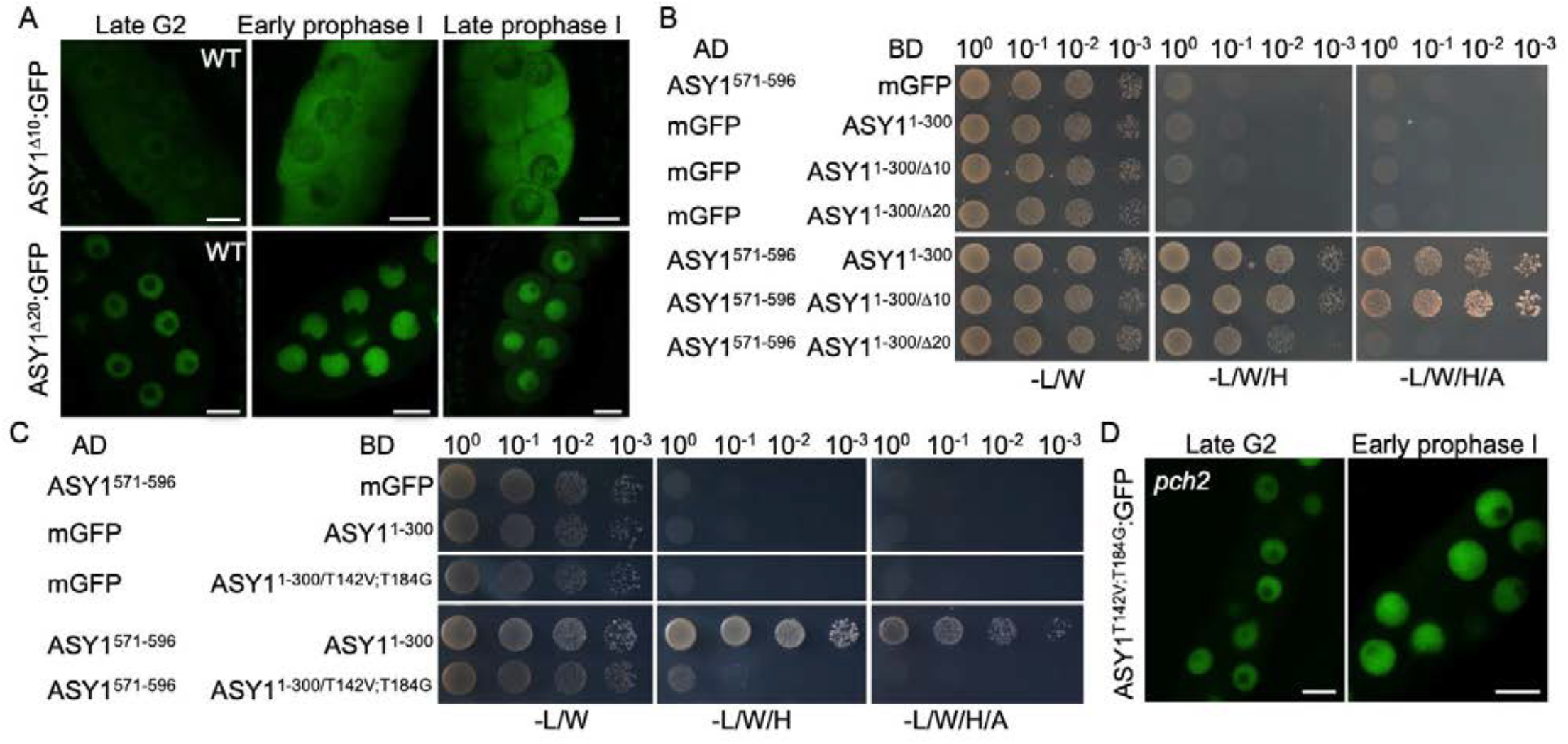
The PCH2-mediated dissociation of the closure motif of ASY1 from the HORMA domain is required for its efficient nuclear targeting. (A) Localization of ASY1^Δ10^:GFP and ASY1^Δ20^:GFP at different meiotic stages in male meiocytes of wildtype (WT) using confocal microscopy. Bar: 10 μm. (B) Yeast two-hybrid assays for the interaction of the closure motif (ASY1^571-596^) with ASY1 HORMA domains having either the first 9 (ASY1^1-300/Δ10^) or 19 (ASY1^1-300/Δ20^) aa deletion at the N-terminus.

If ASY1 in an unlocked state goes into the nucleus more efficiently than in a self-closed state, and if the cytoplasmic retention of ASY1^Δ10^:GFP in the wildtype is attributed to the tight binding of the HORMA domain to the closure motif, a disturbance of this binding should recover the nuclear targeting of ASY1^Δ10^:GFP in the wildtype as well as ASY1:GFP in *pch2* mutants. To test this idea, we generated different versions of ASY1 in which the binding of the HORMA domain to the closure motif is either abolished or largely compromised.

First, considering that the N-terminal HORMA domain (15-228 aa for ASY1) is well-known to function as an intact unit (15, 26, 27), we wondered whether an additional short deletion at the N-terminus of ASY1^Δ10^:GFP would disrupt the interaction of the HORMA domain with the closure motif and thus, recover its nuclear localization in the wild-type background. To test this, we created a version of ASY1 missing the first 2-20 aa (*PRO_ASY1_:ASY1^Δ2-20^:GFP*, called *ASY1^Δ20^:GFP*) (Fig. 1A). Subsequently, yeast two-hybrid assays were performed for checking the binding of the HORMA domain-to-closure motif. Indeed, while ASY1^1-300/Δ10^ showed a strong interaction with the closure motif (ASY1^571-596^), which is indistinguishable from the ASY1^1-300^, the binding of the HORMA domain-to-closure motif was largely compromised when ASY1^1-300/Δ20^ was used for the assay (Fig. 5B). Consistent with this result and the idea that a disturbance of the HORMA domain-to-closure motif binding might recover the nuclear targeting defect of ASY1^Δ10^:GFP in the wildtype, we observed that ASY1^Δ20^:GFP was localized exclusively in the nucleus of wild-type plants at late G2 and early prophase I and only a weak signal in the cytoplasm was detected at late prophase I (Fig. 5A).

Second, based on our previous finding that the mutation of two presumptive CDK phosphorylation sites in ASY1 (T142 and T184) to none-phosphorylatable residues compromises the HORMA domain-to-closure motif binding (18), we generated a new non-phosphorylatable version of ASY1 harboring mutations at these two sites (*PRO_ASY1_:ASY1^T142V;T184G^:GFP*, called *ASY1^T142V;T184G^:GFP*) that almost completely abolishes the interaction of the HORMA domain-to-closure motif as shown by the yeast two-hybrid assay (Fig. 5C). Subsequently, the *ASY1^T142V;T184G^:GFP* construct was introduced into *pch2* mutants (Table 1). Interestingly, we found that ASY1^T142V;T184G^:GFP was mainly localized in the nucleus of male meiocytes in *pch2* mutants and only a weak signal in the cytoplasm was observed (Fig. 5D), lending further support to our hypothesis that ASY1 in an unlocked state can be imported into the nucleus more efficiently.

Taken together, these data suggest that PCH2 facilitates the nuclear targeting of ASY1 by dissociating the binding of the HORMA domain-to-closure motif.

### The position and origin of the closure motif is not important for its functional interplay with PCH2

To test the above-described interplay between the closure motif and PCH2, we replaced the closure motif of ASY1 with the HORMA domain interacting sequence of ASY3 (called closure motif as well) (Fig. 1A). Consistent with previous data, we found that the N-terminus of ASY3 mediates the interaction with the HORMA domain of ASY1 (Fig. S4) (12). Next, we generated the substitution-of-function version of ASY1 that lacked the ASY1 closure motif but contained the HORMA domain binding sequence of ASY3 along with the SV40 NLS to substitute for the nuclear targeting function of the ASY1 closure motif (*PRO_ASY1_:ASY1_1-570_-ASY3^1-50^-NLS:GFP*, called *ASY1^Δclosure^-ASY3^1-50^-NLS:GFP*) (Fig. 1A). This constructs was then transformed into the *pch2* mutants. Remarkably, we found that the nuclear targeting of both ASY1^Δclosure^-ASY3^1-50^-NLS:GFP was dependent on PCH2 in contrast to the PCH2-independent nuclear localization of *ASY1^Δclosure^-NLS:GFP* (Fig. 6A, Table 1).

**Figure 6.**
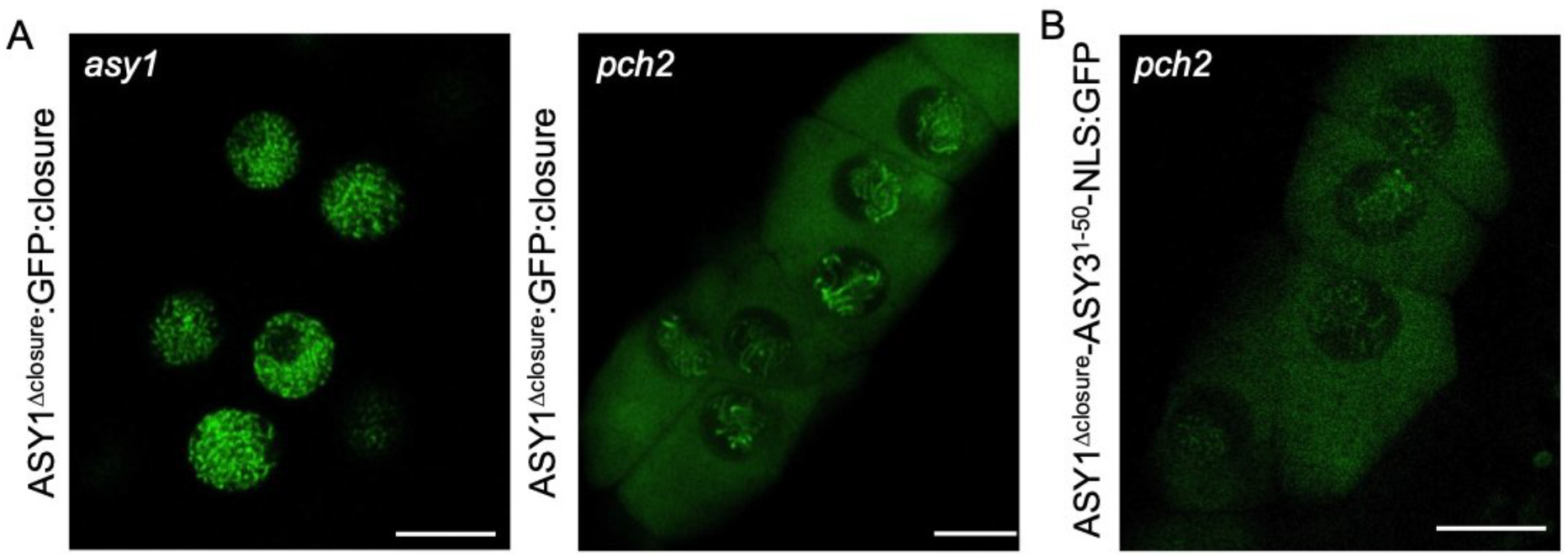
The position and origin of the closure motif is not required for its interplay with PCH2. (A) Localization of ASY1^Δclosure^-ASY3^1-50^-NLS at early prophase in male meiocytes of *pch2* mutants using confocal microscopy. Bar: 10 μm. (B) Localization of ASY1^Δclosure^:GFP:closure at early prophase in male meiocytes of *asy1* and *pch2* mutants using confocal microscopy. Bar: 10 μm.

Finally, we asked to what degree the closure motif-HORMA domain interaction is dependent on the specific context of ASY1. To answer this question, we constructed another version of ASY1 in which the closure motif is separated from the rest of ASY1 by GFP (*PRO_ASY1_:ASY1^1-570^:GFP:ASY1^571-596^*, called *ASY1^Δclosure^:GFP:closure*), which produced an ASY1 variant with an increased distance between the HORMA domain and closure motif (Fig. 1A). This construct was transformed into *asy1* and *pch2* mutants. We found that *asy1* mutants harboring the *ASY1^Δclosure^:GFP:closure* construct were fully fertile resembling *asy1* mutants transformed with the wild-type version of ASY1 (Fig. S1A and B). Moreover, we observed that the nuclear localization of ASY1^Δclosure^:GFP:closure was dependent on the presence of *PCH2* (Fig. 6B, Table 1). These results suggest that the distance between the closure motif and HORMA domain in ASY1 is not important for ASY1 activity and the interaction with PCH2.

Taken together, these results suggest that as long as there is a sequence in ASY1 that binds to its HORMA domain presumably causing ASY1 to adopt a closed state, PCH2 is needed for converting ASY1 into the unlocked state and subsequent nuclear localization.

## Discussion

Meiotic HORMADs including the budding yeast Hop1, mammalian HORMAD1 and HORMAD2, *C. elegans* HORMADs (HTP-1, HTP-2, HTP-3, and HIM3) and Arabidopsis ASY1 play a crucial role in meiotic recombination. Their chromosomal assembly was recently suggested to depend on at least two mechanisms, the initial recruitment by its binding partners such as ASY3 in Arabidopsis, its yeast homolog Red1 and mice homolog SYCP2, and the putative self-assembly (HORMAD oligomerization) mediated by the HORMA domain-to-closure motif interaction (9, 12, 15, 17, 20, 28, 29). While the localization dependency of meiotic HORMADs on its binding partners, e.g. ASY3, has been experimentally proven in many organisms, the self-assembly mechanism of HORMAD proteins has remained poorly characterized (15, 17). The base of the self-oligomerization mechanism is the HORMA domain-to-closure motif interaction. Here, this mechanism has been challenged, at least in Arabidopsis, since ASY1 without the closure motif was found to localize on chromosomes normally (Fig. 1D and S1). In support of this finding, the point mutation K593A in the closure motif of Hop1 largely abolished the interaction of the HORMA domain with the closure motif and had no impact on the chromosomal localization of Hop1 (13, 17).

Instead, our study implicated the closure motif of ASY1 in its chromosomal removal mediated by PCH2, i.e. ASY1 lacking the closure motif was constantly associated with chromosomes and did not dissociate from synapsed chromosomes (Fig. 3B). Since PCH2 acts through the N-terminus of meiotic HORMADs but not the C-terminal closure motif (Fig. 5A) (19), and since ASY1 in an unlocked state is a prerequisite for its binding with ASY3 (9, 15, 18), we propose that the closure motif ensures the success of ASY1 removal from chromosomes through forcing ASY1 in an self-closed state in which the HORMA domain is bound by the closure motif. We propose that this self-closed state prevents the re-binding of the dissociated ASY1 to the chromosome axis via interacting with ASY3 (Fig. 7).

**Figure 7.**
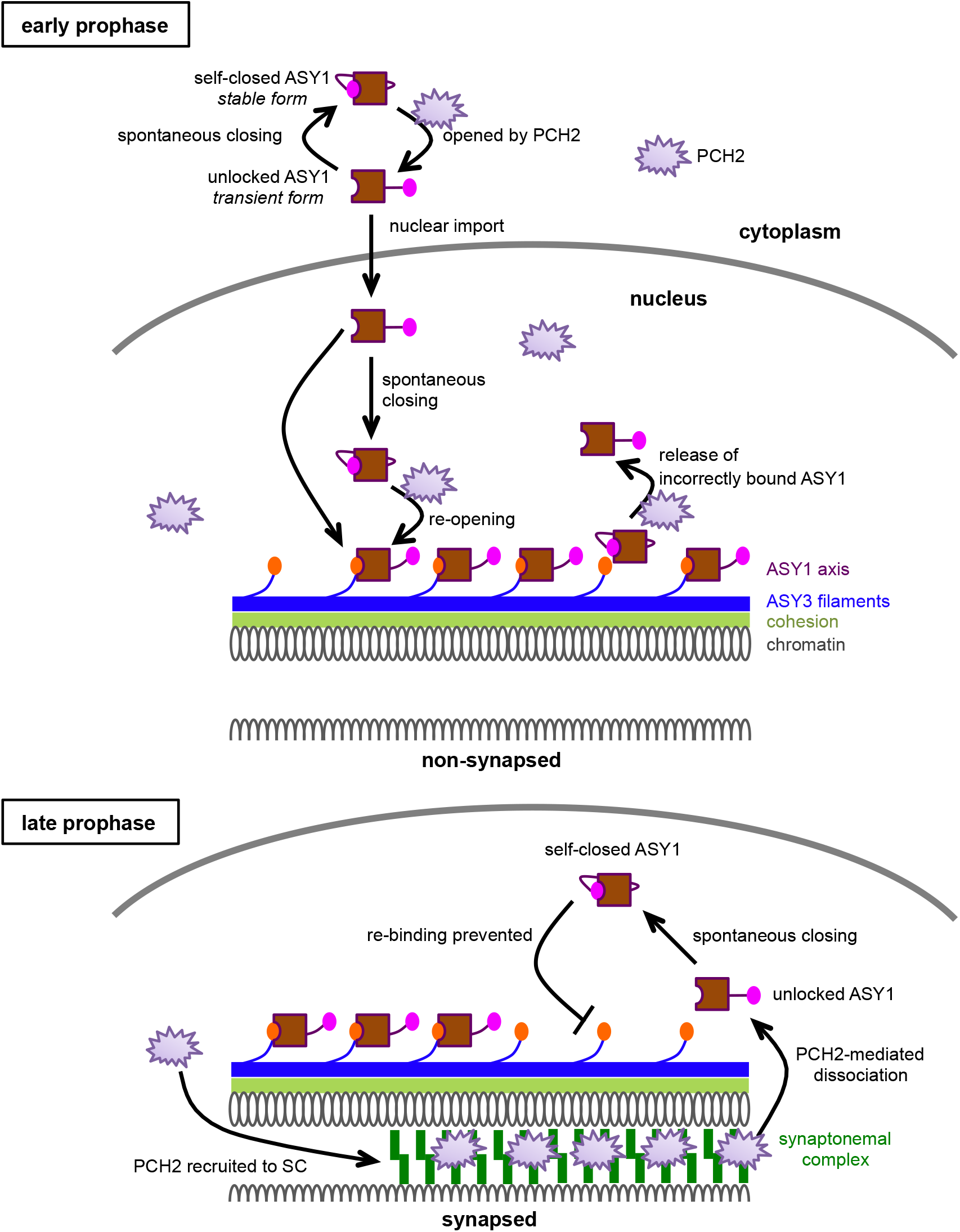
Model of the PCH2-mediated ASY1 dynamics. At early prophase, PCH2 converts the spontaneously self-closing ASY1 to a transient unlocked form by dissociating the closure motif of ASY1 from its HORMA domain allowing the nuclear import of ASY1. PCH2 is diffusely present in the nucleoplasm ensuring the formation of a pool of unlocked ASY1, which is then loaded onto chromosomes via the binding of the HORMA domain to the closure motif of ASY3 giving rise to closed and stably axis-bound form. In addition, PCH2 appears to play a role in a quality control system by releasing unstably/incorrectly bound ASY1 from the chromosome axis (31). Only one homologous chromosome is shown while the second one is indicated by cut DNA loops. At late prophase when homologous chromosomes synapse, PCH2 is recruited to the synaptonemal complex (SC) from the nucleoplasm. The SC-localized PCH2 removes ASY1 from ASY3 by disrupting the interaction of the closure motif of ASY3 with the HORMA domain of ASY1. The so released transient unlocked ASY1 immediately re-folds into a stable, closed form, thus preventing the re-binding of ASY1 to the chromosome axis.

However, this raises the question of why PCH2 does not convert the nucleoplasmic ASY1 depleted from the axis to an unlocked state at late prophase, presumably causing ASY1 to re-bind to the axis by interacting with ASY3. This question is pressing since we reveal here that PCH2 mediates the chromosomal association of ASY1 at early prophase (see below, Fig. 7). The answer to this question possibly lies in the dynamic localization pattern of PCH2. Previous work from us and others show that PCH2 is first diffusedly present in the nucleoplasm of male meiocytes at early prophase when ASY1 needs to be assembled on the axis. In contrast, PCH2 specifically accumulates on synapsed chromosomes and largely decreases in the nucleoplasm when the synaptonemal complex is formed at late prophase (Fig. 7) (18, 21). However, what determines this localization pattern of PCH2 has still to be revealed.

PCH2/TRIP13 proteins have been extensively studied with regard to their function in mediating the dissociation of meiotic HORMADs from chromosomes at late prophase (21, 25, 29, 30). In contrast, their function at early prophase was only recently recognized (31). Here, we show that PCH2 is also indispensable for the proper chromosomal assembly of ASY1 at early prophase in Arabidopsis, as seen by the defective chromosomal association of the ASY1-NLS:GFP proteins in *pch2* mutants but not in the wildtype (Fig. 4C). Interestingly, the dependency of ASY1 localization on PCH2 can be eliminated by deleting the closure motif of ASY1 (Fig. 4B). This suggests a PCH2-mediated conversion of ASY1 from the self-closed state to a transient unlocked state at early prophase, allowing the binding of ASY1 to ASY3 (Fig. 7).

However, it seems that there are at least two pools of ASY1 proteins in Arabidopsis: one pool requires PCH2 for its nuclear targeting and chromosomal association while the other one not. This is evidenced by the strong cytoplasmic retention of some ASY1 molecules in *pch2* mutants versus the clear, albeit weaker than in the wildtype, chromosomal association of some ASY1 proteins (Fig. 4A). Possibly, there is an equilibrium between closed and unlocked ASY1 in which the closed ASY1 is efficiently converted to the unlocked state by PCH2 for nuclear import. When PCH2 is absent, the conformational conversion of ASY1 is blocked and thus, the bulk of ASY1 stays at the closed form failing to enter the nucleus but a little portion in unlocked state that can be imported into nucleus and localizes on chromosomes. However, we cannot fully exclude that there might be other factors regulating/facilitating the nuclear import and chromosomal assembly of the PCH2-independent ASY1 pool. One possibility is that by chance ASY1 and ASY3 may form into a complex in the cytoplasm, which is imported into the nucleus together. However, this remains to be analyzed in the future.

The necessity of PCH2 for both the chromosomal assembly and disassembly of ASY1 during meiosis largely resembles the roles of PCH2/TRIP13 orthologs in the spindle-assembly checkpoint (SAC), a mechanism ensuring that all chromosomes are attached to spindle fibers before anaphase and hence preventing aneuploidy (32–36). PCH2/TRIP13 proteins have been found to be required for not only the inactivation of SAC by disassembling the mitotic checkpoint complex (MCC) comprising BubR1, Bub3, CDC20 and the HORMA protein Mad2, but also for the establishment of the SAC by replenishing the opened Mad2 (O-Mad2) for MCC assembly (33, 37–40). The findings here support a hypothesis that PCH2/TRIP13 proteins promote both the establishment and disassembly of the HORMAD signaling involved in different processes of both mitosis and meiosis in different organisms from yeast to animal and plant species. Our results support the finding that the defects in SAC activation in the absence of PCH2/TRIP13 are likely due to the deficiency of the conformational dynamics of the closed Mad2 (C-Mad2) to O-Mad2, thus abolishing MCC formation (19, 40). Consistent with this assumption, the absence of PCH2/TRIP13 in *C. elegans* and human cells leads specifically Mad2 to an exclusively closed conformation and thus results in the failure of MCC assembly (33, 39). However, since Mad2 does not contain a closure motif, it remains unknown which interacting partners promote Mad2 to a closed state prior to the formation of the MCC.

Our prior study showed that PCH2 in Arabidopsis is essential for the efficient nuclear targeting of ASY1 (18). In this study, by generating a series of ASY1 protein versions the functional interplay between PCH2 and the closure motif with regard to the nuclear targeting and chromosomal localization of ASY1 was revealed (Fig. 4 and 7). We have further demonstrated that PCH2 controls the nuclear targeting of ASY1 through dissociating the interaction between the HORMA domain and the closure motif (Fig. 5). Thus, the unlocked state of ASY1 (the HORMA domain unbound by the closure motif) is a prerequisite for its nuclear import (Fig. 7). Nonetheless, the underlying structural reasons remain to be understood.

The requirement of PCH2 for both the nuclear import and chromosomal assembly of ASY1 might explain the observation that no chromosomal association of PAIR2, the ASY1 homolog in rice, was detected in rice mutants of the *PCH2* ortholog *CRC1* (41). For instance, this could be due to the nuclear targeting defects of PAIR2 and/or the failure of PAIR2 to associate with the chromosome axis when CRC1 is absent. If our finding also holds true in *S. cerevisiae* and mammals, the failure to convert the closed into an unlocked state of Hop1 or HORMAD1/2 in *pch2/TRIP13* mutants could explain the reduced number of DSBs and crossovers in these mutants resembling *hop1* and *homard1/2* mutants (4, 5, 13, 30, 42).

Besides the defective removal from the chromosomes, we reasoned that the inability of ASY1 without closure (ASY1^Δclosure^-NLS:GFP) to complement, not even partially, the fertility and meiotic defects of *asy1* mutants (Fig. S2) might also result from the misregulation of the assembly/loading of components of the recombination machineries. Thus, it seems likely that the closure motif of ASY1 also plays additional roles in meiosis, e.g. promoting crossover formation. This idea is supported by the finding that *hop1* cells harboring Hop1-K593A that localized to chromosomes normally, are defective in crossover formation due to the inability to properly activate the Mek1 kinase (13).

Moreover, we have tested the effects of the origin and position of the closure motif on its interplay with PCH2. Interestingly, we found that regardless of the origin and the distance between the closure motif and HORMA domain, as long as there is a HORMA-domain binding sequence in ASY1 that forces ASY1 to adopt a closed state, PCH2 is required for converting ASY1 into the unlocked state and subsequent the nuclear localization and chromosomal association (Fig. 6). Thus it seems that the presence of the closure motif mainly serves to confer ASY1 a structural plasticity essential for its functions in meiosis (Fig. 7). In this context, only the meiotic HORMADs but not the other types, e.g. Mad2 homologs, harbor the closure motifs in their own protein sequences (15). In addition, the accumulating evidence suggests that the PCH2/TRIP13-dependent conformational dynamics that allow the closure motif binding and dissociation seem to be an evolutionarily conserved mechanism of the HORMA protein signaling. Thus, we speculate that during evolution of the sexual reproduction, especially of meiosis, meiotic HORMADs might either combine the HORMA domain function and their binding regions into one protein sequence, or obtain their own closure motifs through a convergent evolution. Hence, the closure motif likely endows meiotic HORMADs with a more robust regulation with respect to the dynamic chromosomal assembly and disassembly, preventing meiotic defects and ensuring the genome stability over generations. Based on the universal existence of meiotic HORMADs and the closure motifs, together with the functional similarity of PCH2 orthologs, we postulate that the findings revealed here are also conserved in other sexually reproducing organisms.

## Materials and methods

### Plant materials

The Arabidopsis thaliana accession Columbia (Col-0) was used as wild-type reference throughout this research. The T-DNA insertion lines SALK_046272 (*asy1-4*) (43) and SALK_031449 (*pch2-2*) (21) were obtained from the T-DNA mutant collection at the Salk Institute Genomics Analysis Laboratory (SIGnAL, http://signal.salk.edu/cgi-bin/tdnaexpress) via NASC (http://arabidopsis.info/). The *PRO_ASY1_:ASY1:GFP, PRO_ASY1_:ASY1:RFP* and *PRO_ZYP1B_:ZYP1B:GFP* reporters were described previously (18, 22). All plants were grown in growth chambers with a 16 h light/21°C and 8 h/18°C dark cycle at 60% humidity.

### Plasmid construction and plant transformation

To generate the *ASY1^Δclosure^:GFP* reporter, the PCR for the closure motif deletion was performed with the primer pair (gASY1 1-570aa-R and mGFP-F) using the entry clone of *PRO_ASY1_:ASY1:GFP/pENTR2B* generated previously as a template and subsequently the PCR fragments were re-ligated producing the *PRO_ASY1_:ASY1^1-570^:GFP/pENTR2B* construct. For the *ASY1^Δclosure^-NLS:GFP* reporter, the PCR fragments were amplified with the primer pair (gASY1 1-570aa-R+NLS and mGFP) using *PRO_ASY1_:ASY1:GFP/pENTR2B* as a template and then re-ligated, producing the *PRO_ASY1_:ASY1^1-570^-NLS:GFP/pENTR2B* construct. For creating the *Closure:GFP* reporter, the PCR fragments were generated with the primer pair (gASY1-promoterATG-R and gASY1-NLS-571aa-F) using *PRO_ASY1_:ASY1:GFP/pENTR2B* as a template and then re-ligated, producing the *PRO_ASY1_:ASY1^571-596^:GFP/pENTR2B* construct. For creating the *ASY1-NLS:GFP* reporter, the PCR fragments were obtained with the primer pair (gASY1-R+NLS and mGFP-F) using *PRO_ASY1_:ASY1:GFP/pENTR2B* as a template and then religated, producing the *PRO_ASY1_:ASY1-NLS:GFP/pENTR2B* construct. For creating the *ASY1^Δ10^:GFP* and *ASY1^Δ20^:GFP* reporters, PCR fragments were amplified with primer pairs (gASY1-11aa-F and gASY1-intron1-R or ASY1-intron2-F and gASY1-intron1-R) and subsequently self-ligated, generating the *PRO_ASY1_:ASY1^Δ2-10^:GFP/pENTR2B* and *PRO_ASY1_:ASY1^Δ2-20^:GFP/pENTR2B* constructs. For creating the *ASY1^Δclosure^:GFP:Closure* reporter, the backbone of the construct was obtained with primer pairs (ASY1-TGA-F and mGFP-SmaI-R) using *PRO_ASY1_:ASY1^1-570^:GFP/pENTR2B* as a template, and the closure motif insert (ASY1 571-596aa) harboring 30 bp overlapping sequence in both ends with the backbone, was obtained by annealing of two synthesized long primers (ASY1 571-596aa-F and ASY1 571-596aa-R). Next, the *PRO_ASY1_:ASY1^1-570^:GFP:ASY1^571-596^/pENTR2B* construct was generated by integration of the backbone and the insert using a SLICE reaction. For creating the *ASY1^Δclosure^-ASY1^1-50^-NLS:GFP* reporter, two types of PCR fragments with 25bp overlapping sequence in both ends, i.e. ASY1 backbone and ASY3 1-50aa insert, were amplified with two primer pairs (NLS-SmaI-F and gASY1 1-570aa-R, ASY3-SLICE-F and ASY3-SLICE-50aa-R) using the *PRO_ASY1_:ASY1:GFP/pENTR2B* or ASY3/pDONR223 (generated in (22)) as a template. Subsequently, a SLICE reaction were performed by mixing the ASY1 backbone with ASY3 1-50aa fragments, producing the *PRO_ASY1_:ASY1^1-570^-ASY3^1-50^-NLS:GFP/pENTR2B* entry construct. For creating the *ASY1^T142V;T184G^:GFP* reporter, a PCR-based mutagenesis was performed using with primer pair (ASY1 T184G CDS-F and ASY1 T184-R) using the previously generated *PRO_ASY1_:ASY1^T142V^:GFP/pENTR2B* (22) as a template and the resulting fragments were then re-ligated.

Next, all these resulting expression cassettes were integrated into the destination vector *pGWB501/pGWB601* by the gateway LR reaction. All constructs were transformed into *Arabidopsis thaliana* plants by floral dipping.

For constructs of the yeast two-hybrid assays, the *ASY1^571-596^-AD, ASY1^1-300^-BD* and *ASY3-FL-AD* constructs were generated previously (18). To generate *ASY1^1-300/Δ10^-BD* and *ASY1^1-300/Δ20^-BD*, a PCR-based fragment deletion was performed with two primer pairs (ASY1-11aa-F and ATG-attL1-R2, ASY1-21aa-F and ATG-attL1-R2) using *ASY1^1-300^/pDONR223* as a template and the resulting fragments were then re-ligated producing the entry clones. To create *ASY3^1-200^-AD* and *ASY3^201-793^-AD*, the coding sequences of the respective fragments were amplified by PCR with primers flanked by attB recombination sites (ASY3-attB1-F and ASY3-200aa-attB2-R, ASY3-201aa-attB1-F and ASY3-attB2-R) and subcloned into *pDONR223* vector by gateway BP reactions. All these entry clones were subsequently integrated into the *pGADT7-GW* or *pGBKT7-GW* vectors by gateway LR reactions. Primers used for generating all constructs mentioned above are shown in Supplemental Table 1.

### Microscopy

Images of pollen staining were taken using an Axiophot microscope (Zeiss). For the protein localization analyses in male meiocytes, young anthers harboring the relevant reporters were dissected and imaged directly using the Leica TCS SP8 inverted confocal microscope. The meiotic stages for live cell imaging were determined by combining the criteria of the chromosome morphology, nucleolus position, and cell shape (44).

### Yeast two-hybrid assay

Yeast two-hybrid assays were performed according to the manual of Matchmaker Gold Yeast two-hybrid system (Clontech). The relevant combinations of constructs were co-transformed into yeast strain AH109 using the polyethylene glycol/lithium acetate method as described in the manual. Yeast cells expressing the relevant proteins were dotted on the plates of double (-Leu-Trp), triple (-Leu-Trp-His) and quadruple (-Leu-Trp-His-Ade) synthetic dropout medium to test the protein-protein interactions.

### Cytological analysis

Chromosome spread analyses were performed as described previously (18). In brief, fresh flower buds were fixed in fixation solution containing 75% ethanol and 25% acetic acid for 48 h at 4°C. After two times of washing with 75% ethanol, the fixed flower buds were stored in 75% ethanol at 4°C. To perform chromosome spread, flower buds were first digested in the enzyme solution (10 mM citrate buffer containing 1.5% cellulose, 1.5% pectolyase, and 1.5% cytohelicase) for 3 h at 37°C. Subsequently, single flowers were transferred onto a glass slide followed by a fine smashing with a bended needle in the enzyme solution. The spreading step was performed on a 46°C hotplate after adding 10 μl of 45% acetic acid. Subsequently, the slide was rinsed with ice-cold ethanol/acetic acid (3:1) solution and mounted with DAPI solution (Vector Laboratories).

Immunolocalization analyses was performed according to (22). Briefly, fresh flower buds were first sorted by size and then intact anthers likely at meiotic stage were collected and macerated in 10 μl enzyme solution (0.4% cytohelicase and 1% polyvinylpyrrolidone) on the Poly-Prep slides (Sigma) for 5min in a moisture chamber at 37°C followed by a squashing. Next, another 10 μl enzyme solution was added onto the slides that were incubated further for 7 min in the moisture chamber. Subsequently, the anthers were smashed in 20 μl 1% Lipsol for 2 min. Next, 35μl fixation solution (4% (w/v) paraformaldehyde) was added onto the slides followed by a gentle stirring and drying at room temperature for 2-3 h. Afterwards, the slides were washed three times with PBST buffer (PBS with 1% Triton X-100), and were then blocked in PBST containing 1% BSA for 1 h at 37°C in the moisture chamber. Then the slides were incubated with anti-GFP (Takara 632381/JL-8)) (1:100 dilution) and/or anti-ZYP1 (1:500 dilution) antibodies at 4 °C for 48 h. Next, following three times of washing (10 min each) in PBST, the slides were incubated with fluorescein-conjugated secondary antibodies for 24h incubation at 4°C in a moisture chamber. After three times of washing, the DNA was counterstained with antifade DAPI solution (Vector Laboratories). Images were captured using the Leica SP8 laser scanning microscopy.

## Acknowledgements

We thank Dr. Maren Heese (University of Hamburg), Konstantinos Lampou (University of Hamburg), Dr. Kevin D. Corbett (University of California, San Diego) and Dr. Stefan Heckmann (IPK Gatersleben) for the critical reading and constructive comments on this manuscript. This study was supported by core funding of the University of Hamburg and a federal grant from the state of Hamburg (Hybrids – Chances and challenges of new genomic contributions, LFF-FV 36) to SMP and AS.

## Author contributions

C.Y. conceived the research and designed the experiments. C.Y., B.H., S.M.P, and P.C. performed the experiments. C.Y. and A.S. analyzed the data. C.Y. and A.S. wrote the manuscript.

---

**Supplemental Figure 1. The closure motif of ASY1 is not required for its chromosomal localization.** (A) Localization of ASY1:GFP and ASY1^Δclosure^-NLS:GFP at different prophase stages in male meiocytes of *asy1* mutants. Bar: 5 μm. (B) Immunolocalization of ASY1-NLS:GFP and ASY1^Δclosure^-NLS:GFP at different prophase stages in male meiocytes of wild-type (WT) plants. Bar: 5 μm.

**Supplemental Figure 2. Phenotypic analysis of different *ASY1* alleles.** (A) Siliques of wild-type (WT), *asy1* mutant, *ASY1:GFP*(*asy1*), *ASY1^Δclosure^-NLS:GFP(asy1), ASY1-NLS:GFP(asy1), ASY1^Δclosure^-NLS:GFP*(WT), and *ASY1^Δclosure^:GFP:closure(asy1)* plants. (B) Peterson staining for the pollens from the *ASY1* alleles shown in (A). Blue indicates aborted pollen grains. (C) Chromosome spread analysis of *asy1* mutants harboring *ASY1^Δclosure^-NLS:GFP* at different meiotic stages. Bar: 10 μm.

**Supplemental Figure 3. Localization of ASY1:GFP at early prophase in the male meiocytes of *pch2* mutants harboring the wild-type version of *PCH2* or the *PCH2^E283Q^*.** Bar: 10 μm.

**Supplemental Figure 4. Yeast two-hybrid assay for the interaction of ASY1 HORMA domain with different versions of ASY3.** Yeast cells co-transformed with the relevant AD- and BD-tagged proteins were grown on plates of double (-L/W), triple (-L/W/H) and quadruple (-L/W/H/A) dropout medium with different dilutions for 3 days.

