## Supplementary table and figures for "Closure motif-mediated state changes of the HORMA protein ASY1"

#### Supplemental table 1. Primers used in this research.

| Primer name | Sequnence (5' to 3') |
|---|---|
| gASY1 1-570aa-R | CTGTGAGGCTTGGCTACAGTTGACTGTC |
| mGFP-F | CCCGGGGGTGGCatggtgagcaagggcgaggagc |
| gASY1 1-570aa-R+NLS | CACCTTTCTCTTCTTCTTTGGCTGTGAGGCTTGGCTACAGTTGACTGTC |
| gASY1-promoterATG-R | CATttttgcagaagtgtgaaacgaataacgag |
| gASY1-NLS-571aa-F | CCAAAGAAGAAGAGAAAGGTGGACAGACGTGGCAGGAAAACCAGC |
| gASY1-R+NLS | CACCTTTCTCTTCTTCTTTGGATTAGCTTGAGATTTCTGACGCTTGG |
| gASY1-11aa-F | GAGATCACTGAGCAGGACTCGCTTCTTCTGg |
| gASY1-intron1-R | ctggaggtaagaaacaaaacgaaacgctaaaatcagactg |
| ASY1-intron2-F | gtaagctacgccgatcatcgagcttttgagtttttgtttc |
| ASY1-TGA-F | AAATCTCAAGCTAATTGAagacaccacctctatcagacaccataaccacc |
| mGFP-SmaI-R | CCTGCCACGTCTGTCCCCGGGTCCACCTCCcttgtacagctcgtccatg |
| ASY1 571-596aa-F | GGAGGTGGACCCGGGGACAGACGTGGCAGGAAAACCAGCATGGTGAGGGAGCCTATTCTGCAGTACT<br>CCAAGCGTCAGAAATCTCAAGCTAATTGAagacaccacctc |
| ASY1 571-596aa-R | gaggtggtgtctTCAATTAGCTTGAGATTTCTGACGCTTGGAGTACTGCAGAATAGGCTCCCTCACCATG<br>CTGGTTTTCCTGCCACGTCTGTCCCCGGGTCCACCTCC |
| NLS-SmaI-F | CCAAAGAAGAAGAGAAAGGTGCCCGGGGGTGGC |
| ASY3-SLICE-F | AGTCAACTGTAGCCAAGCCTCACAGATGAGCGACTATAGAAGCTTCGGCA |
| ASY3-SLICE-50aa-R | CGGGCACCTTTCTCTTCTTCTTTGGCAACTTTTCTACTCTAGCAATAAC |
| ASY3-SLICE-100aa-R | CGGGCACCTTTCTCTTCTTCTTTGGCTCAAGAGTCCCTAATTTCCGATG |
| ASY1 T184G CDS-F | GGGCCACCAGATTACGAGCCACCTT |
| ASY1 T184-R | CACATCATCGTAGTACAGAAGCTTC |
| ASY1-11aa-F | GAGATCACTGAGCAGGACTCGCTTCTTCTG |
| ATG-attL1-R2 | CATGAAGCCTGCTTTTTTGTACAAAGTTGG |
| ASY1-21aa-F | ACTAGAAATTTGCTTCGTATTGCTATCTTC |
| ASY3-attB1-F | GGGGACAAGTTTGTACAAAAAAGCAGGCTTAATGAGCGACTATAGAAGCTTCGGCA |
| ASY3-200aa-attB2-R | GGGGACCACTTTGTACAAGAAAGCTGGGTTTCACAGTTTTGATCTCAAAACATCAGT |
| ASY3-201aa-attB1-F | GGGGACAAGTTTGTACAAAAAAGCAGGCTTCTGGGAGATATTGGGAAAAGCTTCAC |
| ASY3-attB2-R | GGGGACCACTTTGTACAAGAAAGCTGGGTTATCATCCCTCAAACATTCTGCGACA |

Supplemental Figure 1

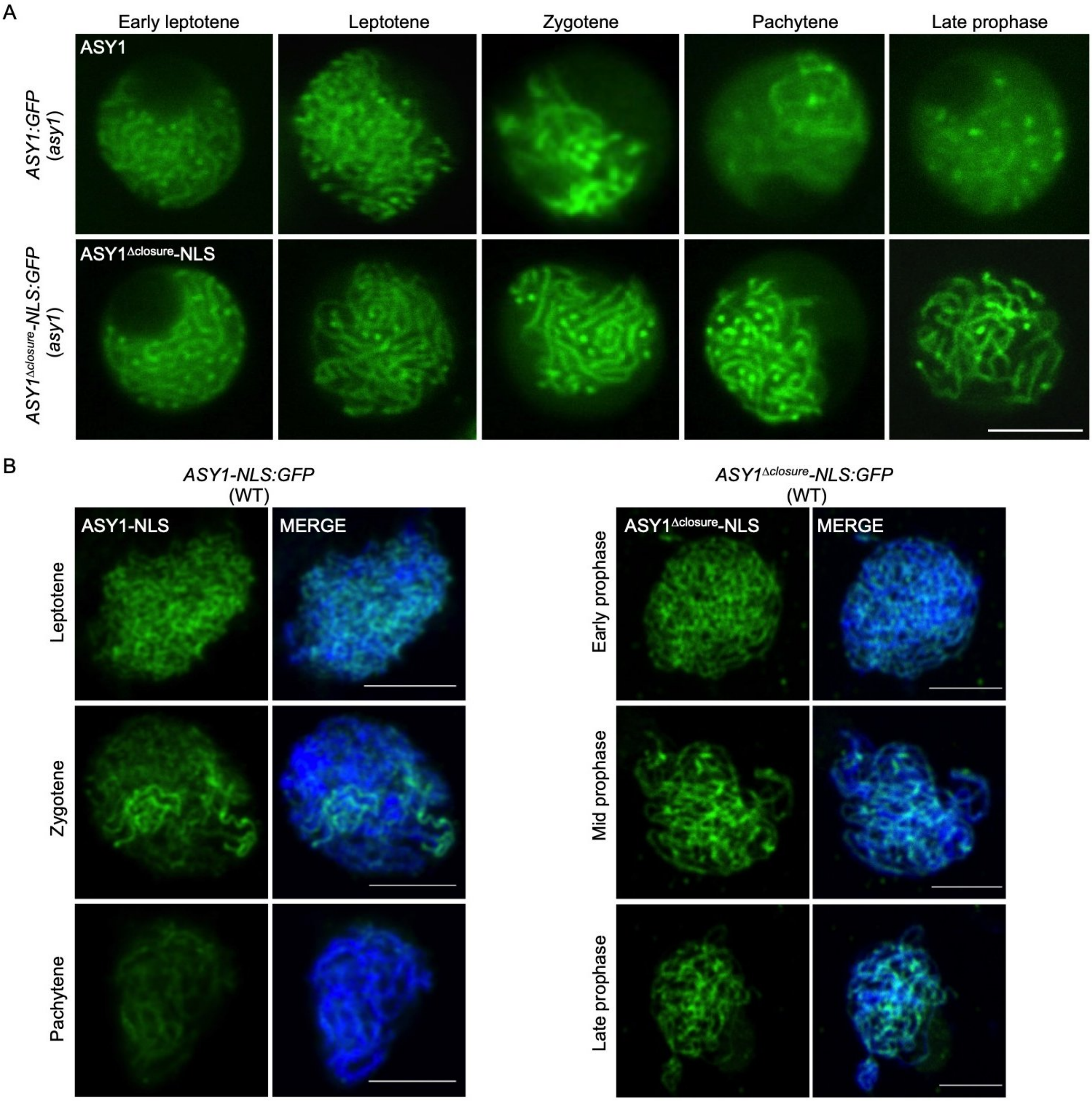

### Supplemental Figure 2

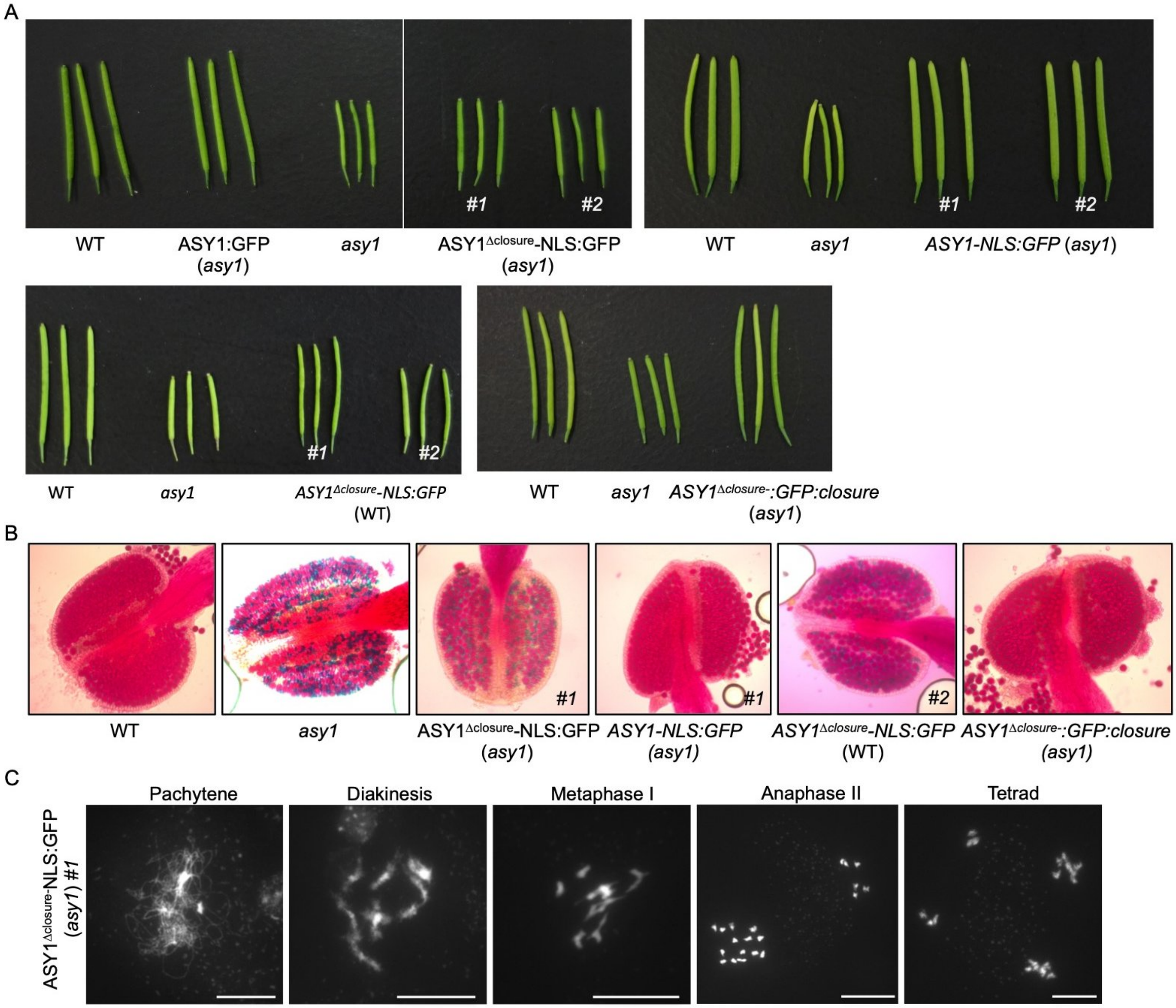

#### Supplemental Figure 3

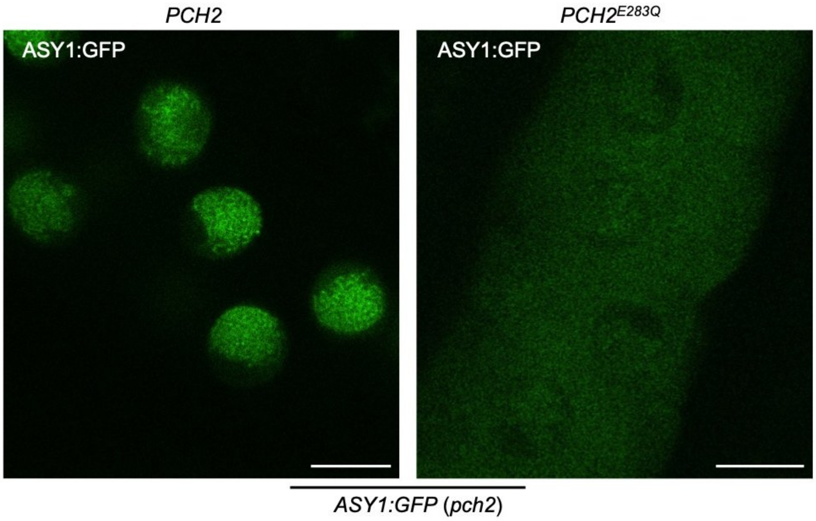


#### Supplemental Figure 4

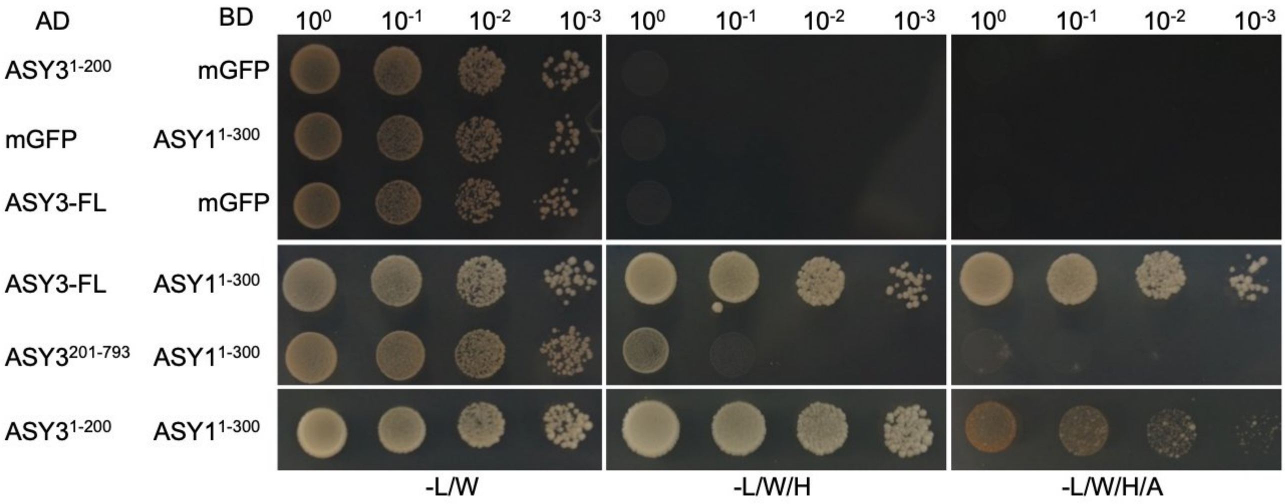
